# The rasterdiv package for measuring diversity from space

**DOI:** 10.1101/2024.02.07.579266

**Authors:** Matteo Marcantonio, Elisa Thouverau, Duccio Rocchini

**Affiliations:** Earth & Life Institute, University of Louvain (UCLouvain), Croix du sud 4-5, 1348 Louvain-la-Neuve, Belgium; BIOME Lab, Department of Biological, Geological and Environmental Sciences, Alma Mater Studiorum University of Bologna, Bologna, Italy; Name, BIOME Lab, Department of Biological, Geological and Environmental Sciences, Alma Mater Studiorum University of Bologna, Bologna, Italy

## Abstract

This book chapter provides an extensive overview of the *rasterdiv* package which provides a comprehensive suite of functions designed to calculate diversity indices from numerical matrices, including those derived from optical remote sensing imagery. It facilitates a deeper analysis of spatial heterogeneity and ecosystem complexity by translating pixel values into ecological indicators. This package serves as a valuable tool for researchers and practitioners in the fields of landscape ecology, biodiversity monitoring, and environmental management, allowing for an enhanced understanding of landscape patterns and their ecological implications through advanced quantitative measures.

## 1 Ecological indexes of ecosystem complexity

Landscape heterogeneity, indicative of land cover diversity, is a pivotal concept in both remote sensing and ecological studies [7]. This diversity is traditionally quantified through algorithms like Shannon’s, Gini-Simpson’s, and Berger-Parker’s indices. These metrics, based on spectral classes (e.g., pixels with the same digital value) within remote sensing data, serve as point-based descriptors of diversity [12] (refer to Table 1). Renyi’s entropy index unifies these indices into a single formula, offering the flexibility to transition between them by adjusting one parameter [7].

**Table 1.**
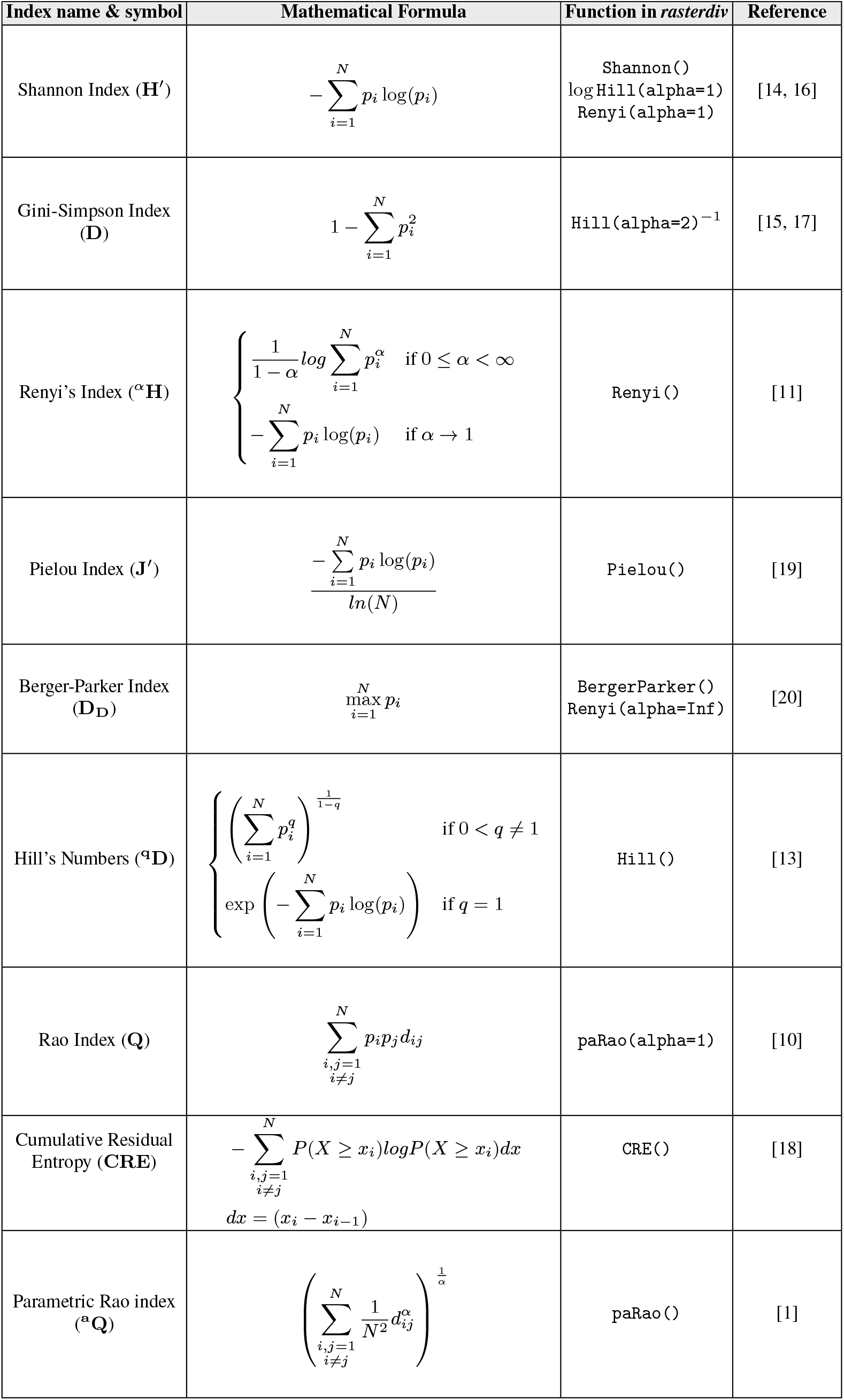
Diversity indices in *rasterdiv* with their mathematical formulas and corresponding functions and references listed in temporal order of first publication.

| Index name & symbol | Mathematical Formula | Function in *rasterdiv* | Reference |
| --- | --- | --- | --- |
| Shannon Index ($\mathbf{H}'$) | $-\sum_{i=1}^N p_i \log(p_i)$ | Shannon()<br>log Hill(alpha=1)<br>Renyi(alpha=1) | [14, 16] |
| Gini-Simpson Index ($\mathbf{D}$) | $1 - \sum_{i=1}^N p_i^2$ | Hill(alpha=2)<sup>-1</sup> | [15, 17] |
| Renyi's Index (${}^{\alpha}\mathbf{H}$) | $\begin{cases} \frac{1}{1-\alpha} \log \sum_{i=1}^N p_i^{\alpha} & \text{if } 0 \leq \alpha < \infty \\ -\sum_{i=1}^N p_i \log(p_i) & \text{if } \alpha \rightarrow 1 \end{cases}$ | Renyi() | [11] |
| Pielou Index ($\mathbf{J}'$) | $\frac{-\sum_{i=1}^N p_i \log(p_i)}{\ln(N)}$ | Pielou() | [19] |
| Berger-Parker Index ($\mathbf{D_D}$) | $\max_{i=1}^N p_i$ | BergerParker()<br>Renyi(alpha=Inf) | [20] |
| Hill's Numbers (${}^q\mathbf{D}$) | $\begin{cases} \left( \sum_{i=1}^N p_i^q \right)^{\frac{1}{1-q}} & \text{if } 0 < q \neq 1 \\ \exp \left( -\sum_{i=1}^N p_i \log(p_i) \right) & \text{if } q = 1 \end{cases}$ | Hill() | [13] |
| Rao Index ($\mathbf{Q}$) | $\sum_{\substack{i,j=1 \\ i \neq j}}^N p_i p_j d_{ij}$ | paRao(alpha=1) | [10] |
| Cumulative Residual Entropy ($\mathbf{CRE}$) | $-\sum_{\substack{i,j=1 \\ i \neq j}}^N P(X \geq x_i) \log P(X \geq x_i) dx$<br>$dx = (x_i - x_{i-1})$ | CRE() | [18] |
| Parametric Rao index (${}^a\mathbf{Q}$) | $\left( \sum_{\substack{i,j=1 \\ i \neq j}}^N \frac{1}{N^2} d_{ij}^{\alpha} \right)^{\frac{1}{\alpha}}$ | paRao() | [1] |

Additionally, Rao’s quadratic entropy (referred to as Rao’s index [10]) introduces more complex measures. It incorporates ‘spectral’ distances among pixel reflectance values, thereby enriching the interpretation of diversity [8]. Rao’s index’s versatility is further enhanced through parameterisation, affecting the emphasis on spectral distances [1].

The *rasterdiv* R package provides an array of user-friendly functions for calculating these indices, enabling the assessment of land cover heterogeneity from satellite imagery. *rasterdiv* employs abundance-based metrics to transform spectral data from matrix-like objects (e.g., *terra*’s *SpatRaster*) into layers representing various diversity indices. This approach considers neighbouring cell values, thus offering a more nuanced local diversity estimation (see Table 1).

In this chapter, we explore the practical applications of the *rasterdiv* package using two distinct datasets, each offering different ecological insights tied to the reported examples.

The first dataset comprises three Normalized Difference Vegetation Index (NDVI) raster layers at a spatial resolution of 60 cm sourced from the National Agriculture Imagery Program (NAIP). These layers represent the southern area of the Berryessa Snow Mountain National Monument in California (USA), captured on the 25th, 28th and 30th of May 2016, 2020 and 2022, respectively. This site was selected for its remarkable environmental diversity within a compact area, from riparian woodland to chaparall and grassland plant communities (Figure 1 2012). Moreover, it has recently undergone significant ecological changes following the destructive Wragg wildfire in the summer of 2015 and LNU Complex Fires in August 2020, making it an ideal subject for studying post-disturbance recovery and landscape temporal heterogeneity (Figure 1 2022).

**Fig. 1.**
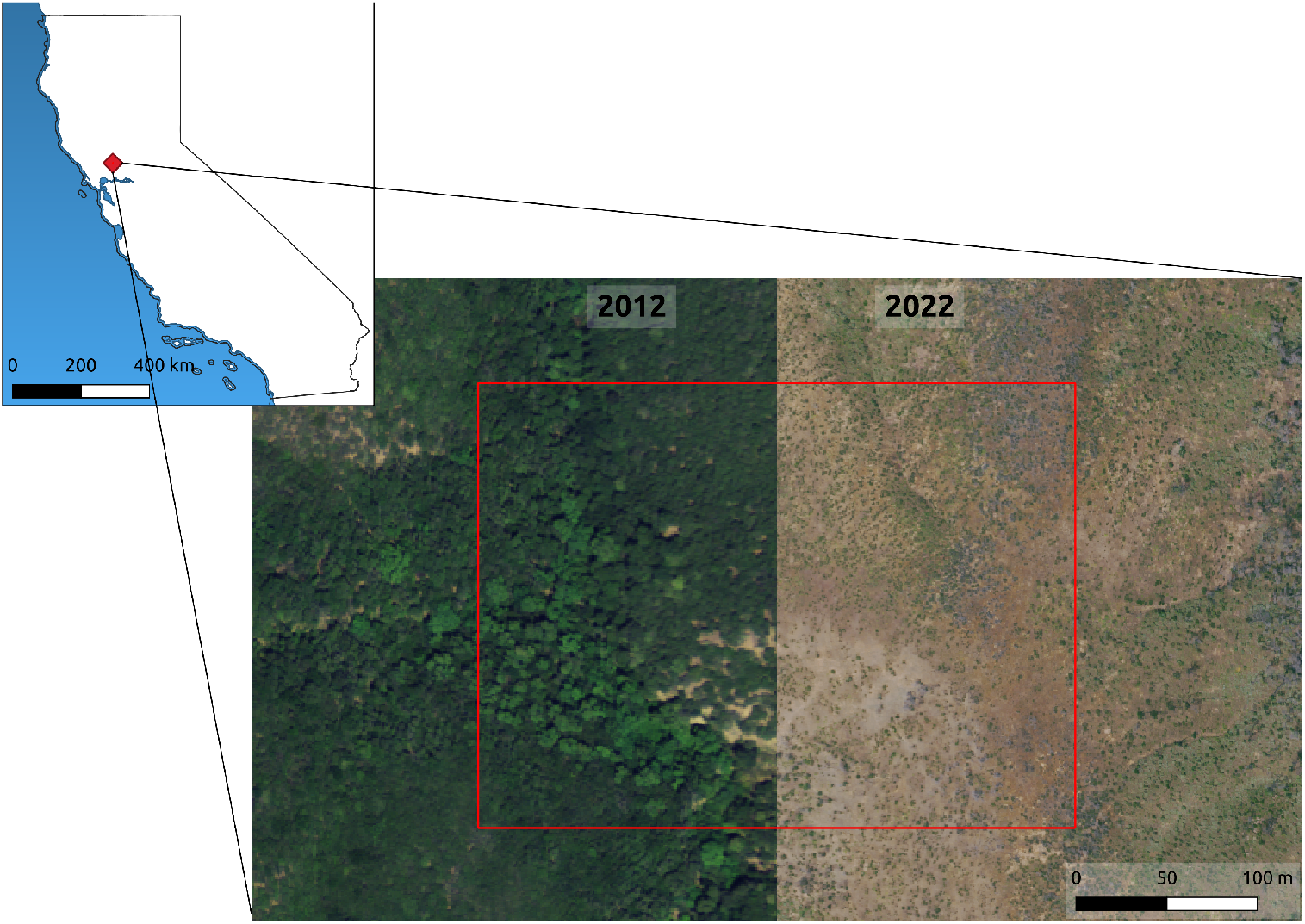
A portion of the UC Davis Stebbins Cold Canyon Reserve, Calufornia, USA (lat: 38.489, lon: -122.101), pre- and post-wildfires in 2012 and 2022, respectively. The red square depicts the case study area used in the first part of this chapter.

The second dataset we will explore is a multi-year (2018-2021) time series of weekly NDVI data from a 20 hectares segment of the Macchiarvana forest in the Abruzzo, Lazio and Molise National Park, Italy derived from 3-meters spatial resolution reflectance values from 4-band PlanetScope PS2 imagery [9]. This forest is predominantly an old-growth beech trees (*Fagus sylvatica*) ecosystem, known for its relatively consistent vegetative dynamics over the years (Figure 2). The expectation here is that the year-on-year NDVI values will display a regular sinusoidal pattern, with minimal variations. Such stability in vegetative growth patterns provides an excellent opportunity to apply *rasterdiv* helical plot functions [3] in understanding the nuances of ecological consistency and the subtle changes that might occur in an old-growth forest environment [4].

**Fig. 2.**
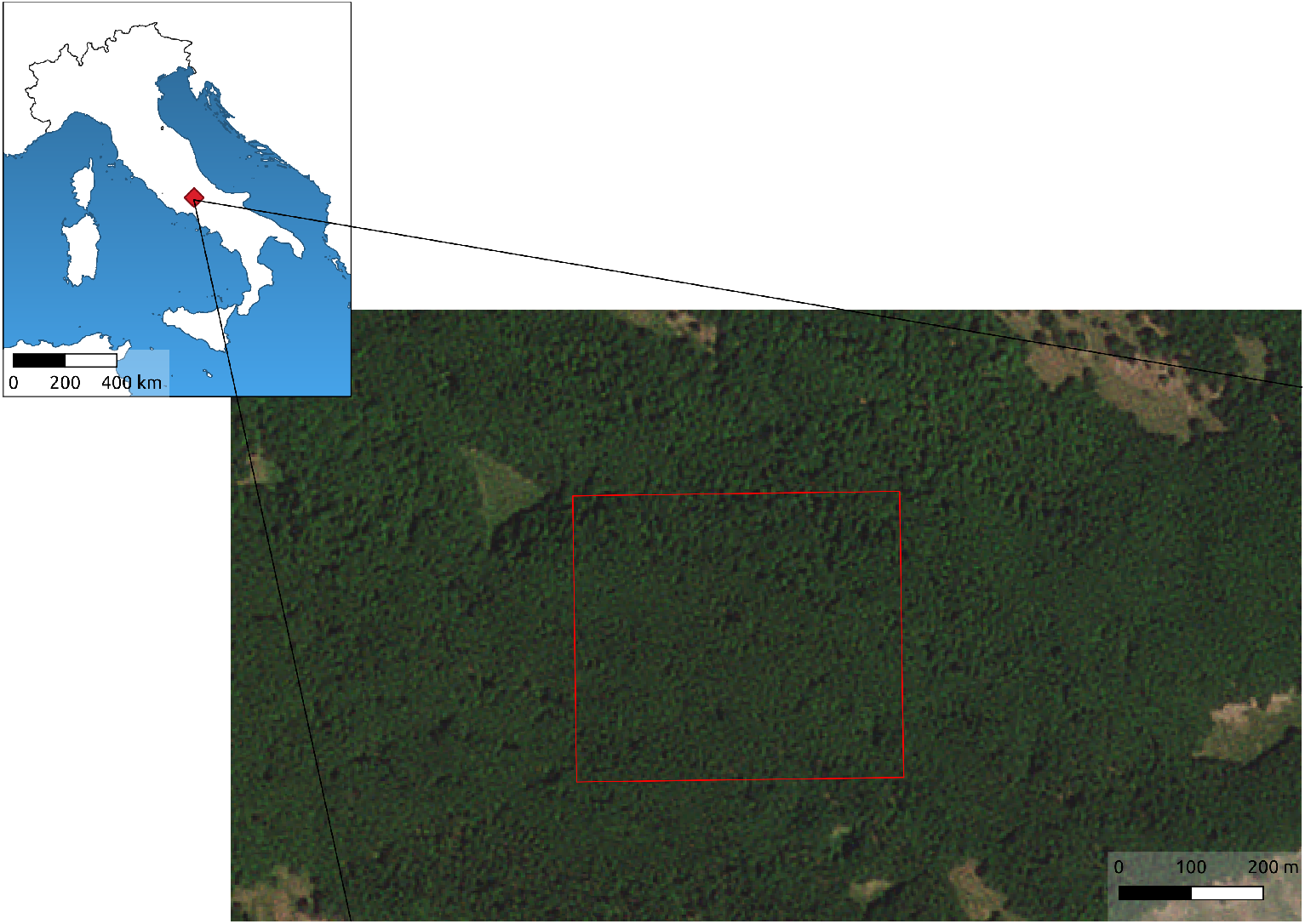
A segment of the Macchiarvana old-growth beech forest located in the Central Appennins, Abruzzo, Italy (lat: 41.766, lon:13.799). The red square depicts the case study area used in the second part of this chapter.

Through these datasets, we aim to illustrate the diverse applications of *rasterdiv* in assessing and interpreting ecological dynamics and landscape heterogeneity, utilising NDVI as a key indicator of spatial heterogeneity in vegetative health and activity. The two datasets used in this chapter can be accessed through the Open Science Framework’s permanent repository. For more information and to download the datasets, please visit the following link: *rasterdiv* datasets on Open Science Framework https://osf.io/pgbmq/.

## 2 Classical indexes of diversity in the *rasterdiv* R package

### 2.1 Preparing raster layers

To effectively apply the diversity indices provided by the *rasterdiv* package, it is imperative to begin with raster layers composed of a finite number of discrete values. Typically, these are represented as 8-bit integers ranging from 0 to 255. The rationale for this requirement lies in the fact that the indices in *rasterdiv* are designed to handle discrete data categories—such as integer counts of spectral classes or land cover types—rather than continuous data.

The indices calculate various transformations based on the relative abundance of pixel values, interpreted as distinct “types” (e.g., spectral classes) within the raster. When the raster data contains a vast range of continuous numeric values, particularly with high precision such as three decimal places, the resulting high number of unique classes coupled with an extremely low relative abundance per class can render the diversity calculations meaningless. In such cases, the diversity indices may not provide insightful information due to the dilution of meaningful class distinctions.

**Essential Tips for Utilising *rasterdiv***

- **Input Data [*x*]**: Essential to all analyses, input data should be a numerical matrix or a *SpatRaster* layer with discrete cell values. Typically, an 8-bit numeric format is preferred to ensure accurate computation of diversity indices.
- **Moving Window [*window*]**: This square kernel, central to the analysis, traverses the image pixel by pixel. Its size is crucial as it determines the scope of the local neighborhood considered for each pixel during index calculations.
- **NA Tolerance [*na*.*tolerance*]**: This parameter sets the proportional limit for missing data within each moving window, crucial for handling boundaries and data gaps accurately. Exceeding this limit results in an NA value for the central pixel’s index, ensuring integrity in areas where the dataset is incomplete.
- **Parallel Processing [*np*]**: To enhance computational efficiency, the package allows for parallel processing. The ‘number of processes’ parameter
- enables users to specify the number of concurrent processing threads, thereby optimising performance on multi-core systems.
- **Value Simplification [*simplify*]**: In dealing with high-precision floating-point numbers, simplifying pixel values by rounding them to a specified decimal place can significantly streamline index computation, balancing accuracy with processing demands.

Therefore, for the *rasterdiv* functions to yield significant and interpretable results, preprocessing steps may be necessary to categorise continuous raster data into a manageable number of discrete classes that reflect meaningful differences in the landscape or the spectral characteristics being studied. The linear transformation from floats to 8-bit (or any other range of numeric values) can be achieved using *terra::stretch()* function, by setting the output range of values between 0 and 255.

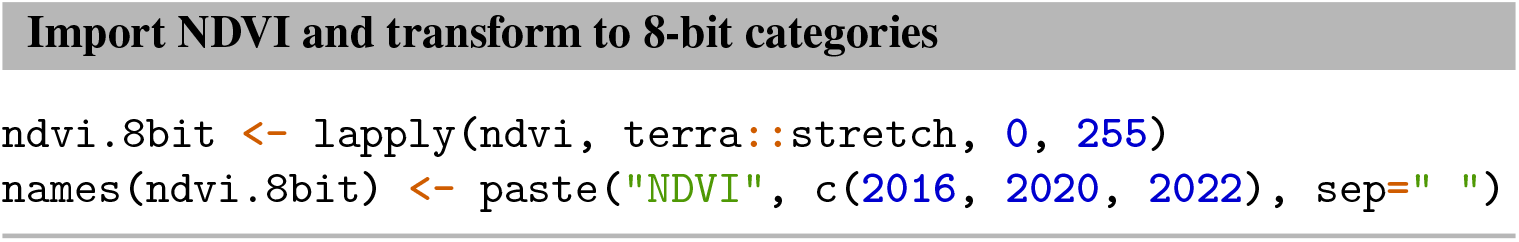

Throughout this chapter, we demonstrate the use of *rasterdiv* functions alongside *lapply*, a strategy that effectively iterates over multiple *SpatRasters*.

### 2.2 Renyi’s generalised entropy

Renyi’s entropy [11], a generalisation of the Shannon entropy formula denoted ^*α*^*H*, offers a versatile approach to assess the prominence of different spectral classes in a landscape (Table 1). By adjusting the *α* parameter, one can effectively modulate the emphasis placed on the most dominant spectral class. This adaptability is integrated into *rasterdiv* via the *alpha* option in the *Renyi()* function. The *Renyi()* function in *rasterdiv*, along with all other diversity index functions, is designed to handle large datasets by leveraging parallel computing capabilities. Users can optimise performance through the *cluster*.*type* option, which supports classical parallel computation backends including “SOCK”, “FORK”, and “MPI”. Additionally, the degree of parallelism (which acts at single matrix layers) can be customised by specifying the number of processes using the *np* option. This feature significantly enhances the processing speed, making it possible to analyse extensive spatial data effectively.

To demonstrate the utility of Renyi’s entropy index in ecological analysis, we applied it with *α* values of 0, 1, and 2 to three NDVI datasets (embedded in the *ndvi*.*8bit* multilayer *SpatRaster*) of Stebbins Cold Canyon (Figure 3). These datasets correspond to three different ecological moments: immediately following the Wragg wildfire in 2016, just prior to the LNU wildfire in 2020, and two years post the LNU wildfire in 2022.

**Fig. 3.**
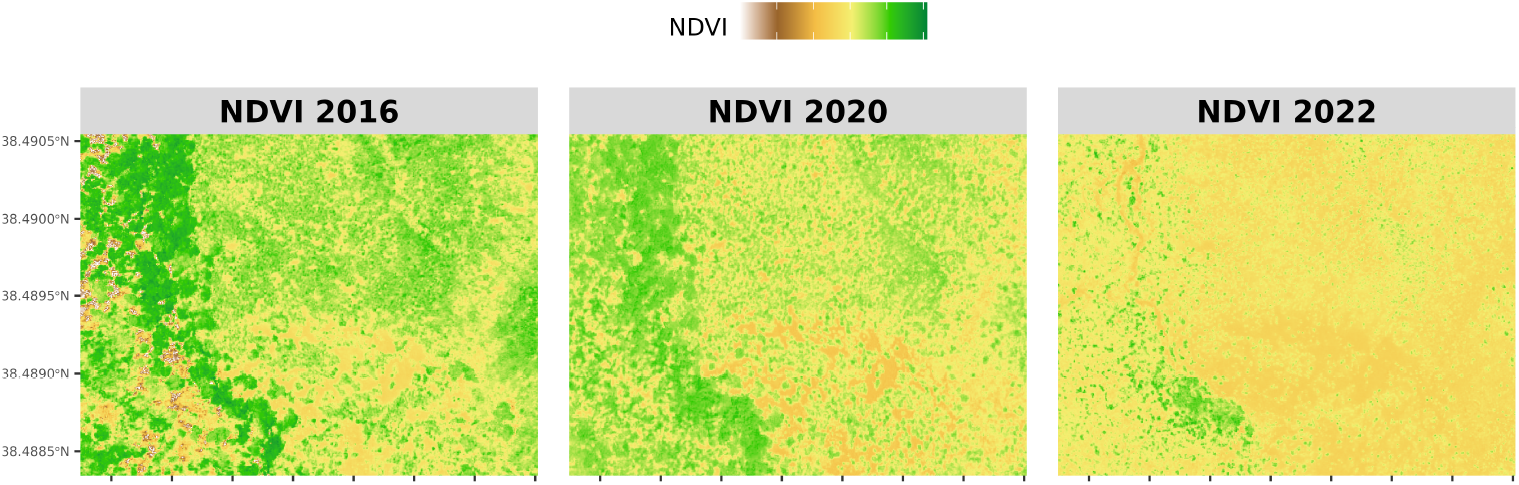
NDVI values of the study areas in three different years. Higher NDVI values indicate wetter river plant communities at the bottom of the canyon, whereas lighter values chapparal and grassland communities at the top of the hill ridge.

This application allows us to explore the heterogeneity and ecological dynamics of the landscape under varying degrees of disturbance and recovery. By adjusting the *α* parameter, we can shift our focus from the general diversity (*α* = 0) to the more nuanced aspects of species dominance (higher *α* values), thereby offering a comprehensive view of the landscape’s ecological resilience and response to wildfires over time.

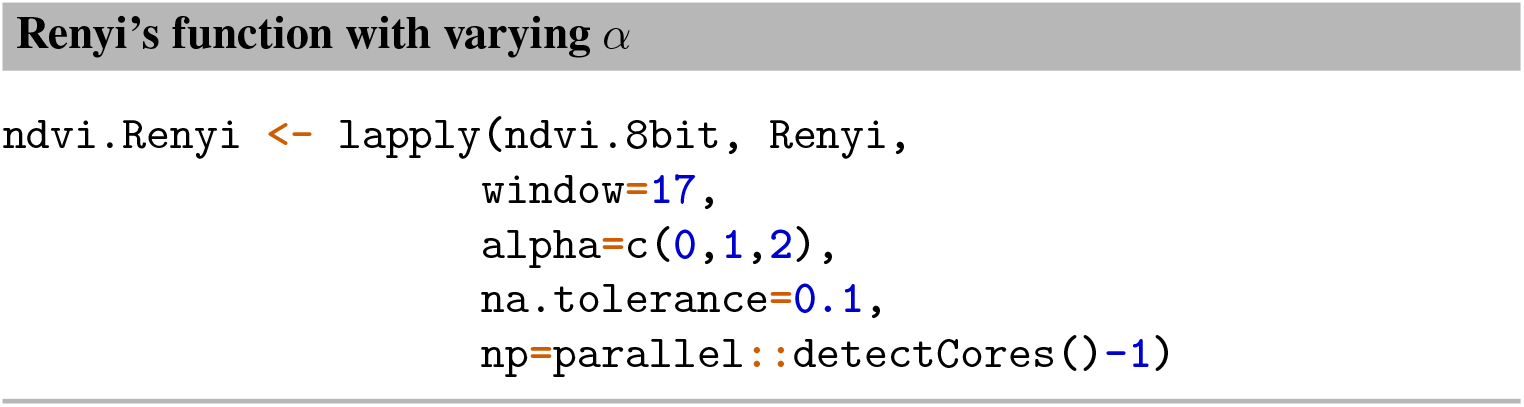

The *ndvi*.*Renyi* list, generated from the application of the Renyi formula across different NDVI datasets, reflects variations in *α* values. Each sublist within *ndvi*.*Renyi* is a lower level list corresponding to a specific moving window value, then each element in the window sublist is a *SpatRaster* (or matrix) object corresponding to each *α* value:

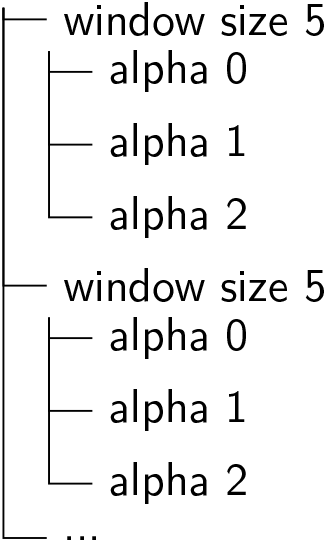

Such hierarchical organisation is common for the outputs of all the diversity index functions provided in *rasterdiv*. This structure facilitates conversion into a multilayer SpatRaster by using *lapply* in conjunction with *terra::rast()*, enabling visualisation through *ggplot2* and *tidyterra* (Figure 4).

**Fig. 4.**
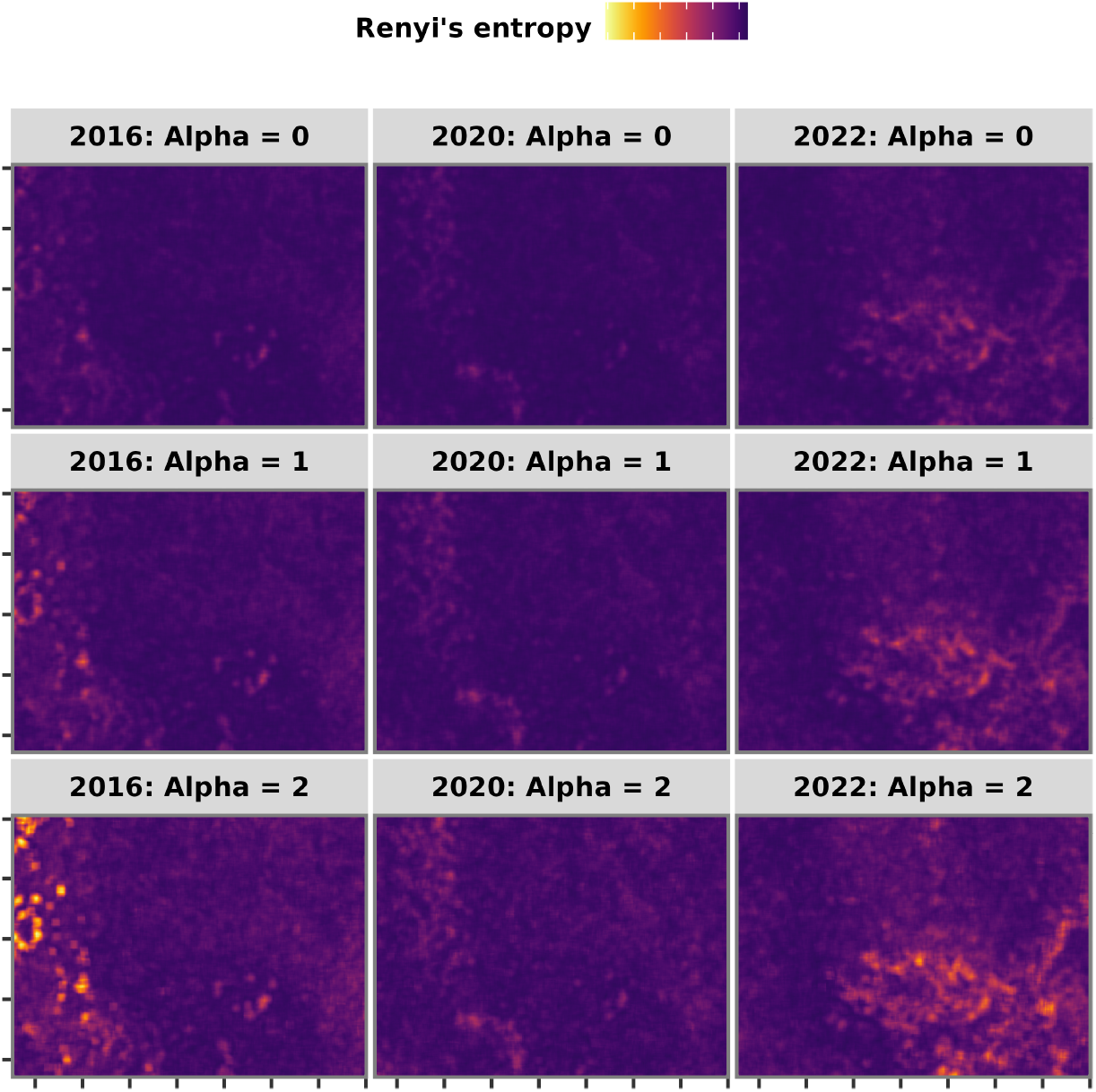
Overview of Renyi’s index with varying *α* values (rows) for NDVI in different years (columns). Lighter colours show lower Renyi’s index values which generally indicate more homogenous habitats.

**Fig. 5.**
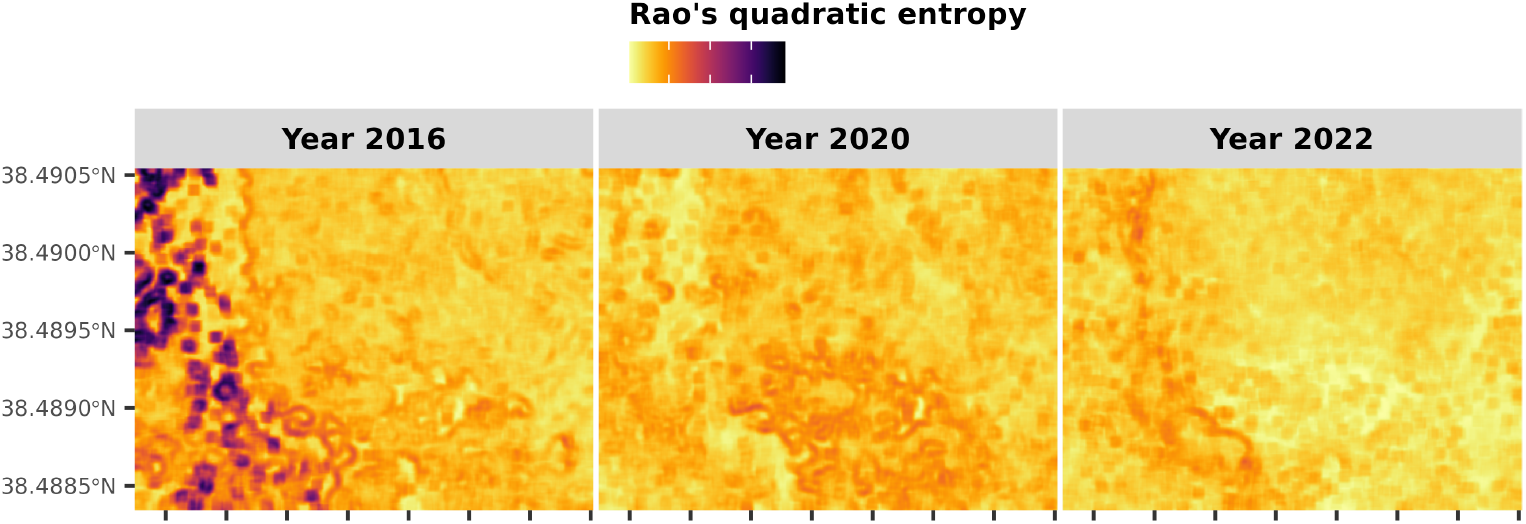
Overview of parametric Rao’s index for NDVI in different years. Darker values show areas where pixel distances and thus spectral heterogeneity is higher. In 2016, areas with portion having freshly burned vegetation are highlighted due to the strong pixel distance between green healthy and burned vegetation patches.

In *rasterdiv*, the *window* parameter in diversity index functions specifies the size in pixels of the square kernel (often referred to as the “moving window”) utilized to compute the relative abundances of spectral classes (see Info Box 2.1). It is crucial that *window* is an odd integer, as it represents a square kernel’s dimensions. While varying *α* offers insights into the dominance of spectral classes, altering the window size can reveal how local landscape heterogeneity influences diversity metrics. Varying the window size in *Renyi()* and all other diversity function contained in *rasterdiv* can be achieved using a vector of odd integers as follows:

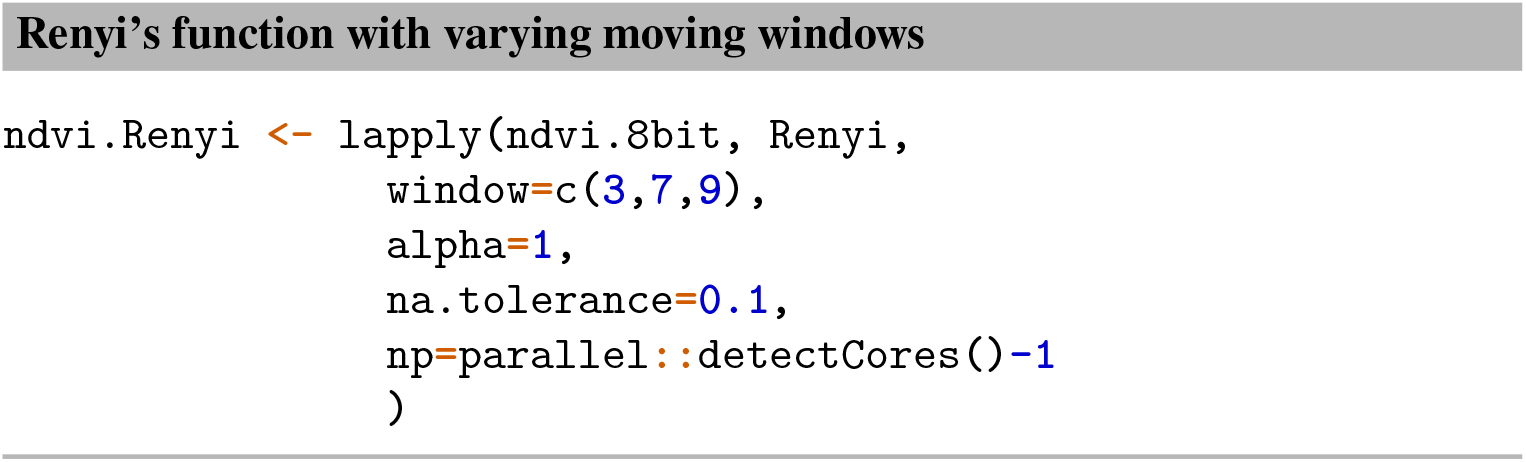

This approach enables ecologists and remote sensing experts to tailor the analysis to specific spatial resolutions, providing a flexible tool for investigating ecological patterns and processes at multiple scales.

### 2.3 Other classical index of diversity

Hill’s numbers, denoted as ^*q*^ *D* and also known as true diversity indices, are a versatile set of measures in biodiversity studies (Table 1). These indices can be effectively computed using the *Hill()* function in the *rasterdiv* package by specifying various *α* values. This functionality parallels that of the *Renyi()* function, offering a broad spectrum of diversity measurements based on different *α* parameters. Beyond these generalised formulas, *rasterdiv* also includes functions dedicated to calculating specific point diversity indices. For instance, the *Shannon()* and *BergerParker()* functions provide a direct approach to computing Renyi’s index for *α* values of 1 and inf, respectively (Table 1).

Additionally, *rasterdiv* offers the *Pielou()* function (*J*′), designed to calculate Pielou’s evenness index (Table 1). This index represents Shannon’s entropy normalised by its maximum possible value, thus ranging between 0 and 1. Pielou’s evenness index is particularly valuable for comparative analyses across different ecological scenarios or temporal scales, as it provides a standardised measure of species distribution uniformity within a given area.

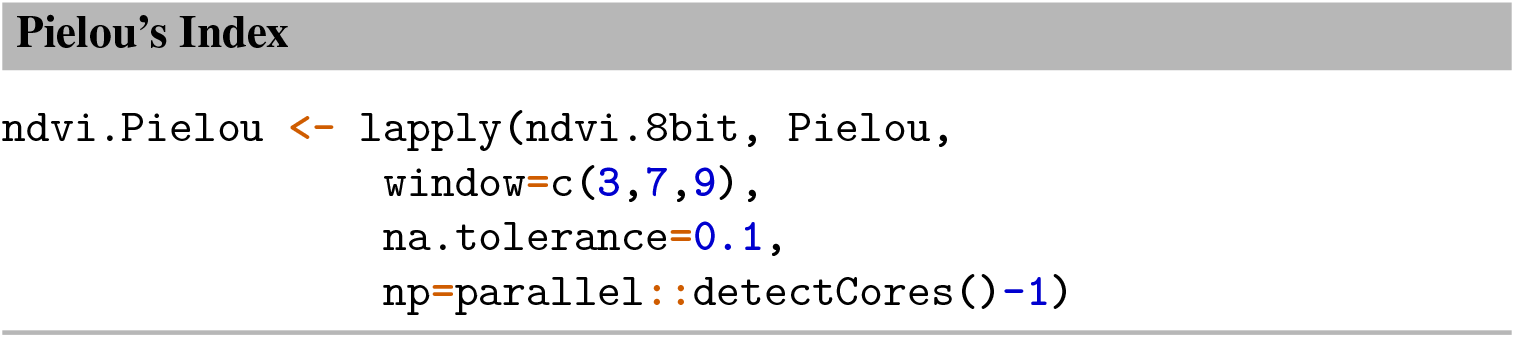

*Pielou()* and all the function reported in this paragraph allows for parameters relevant for their calculation as in the *Renyi()* function.

## 3 Incorporating Distance into Diversity Assessment: Rao’s Quadratic Entropy

In the realm of landscape heterogeneity analysis, diversity indexes are pivotal tools. They often involve various transformations of the relative abundances of pixel ‘types’ (e.g., spectral classes) within a defined spatial context, a moving window in *rasterdiv*. Among these indexes, Rao’s Quadratic Entropy stands out for its ability to integrate ‘distances’ between spectral classes into the diversity index calculation. This more nuanced measure considers not just the presence or abundance of spectral classes but also how different they are from each other in a given characteristic, adding a layer of ecological or spectral interpretation to the analysis.

In the *rasterdiv* package, this index is accessible through the *paRao()* function. This function implements a generalised version of Rao’s formula, as detailed in [1]’s work. Much like the Renyi’s and Hill’s formulas, Rao’s parametrised quadratic entropy formula allows for the adjustment of the index’s emphasis through the *alpha* parameter. In this context, the *alpha* parameter plays a critical role – the higher its value, the greater the emphasis on the distances between spectral classes. When *α* is set to 1, the function reverts to computing the classic Rao’s index, which considers only the distances without additional weighting, thus providing a baseline measure of diversity that reflects the heterogeneity in pixel classes based on their spectral distances. The *paRao()* accommodates a variety of distance metrics to suit diverse analytical needs. Supported distance measures include commonly used options such as “euclidean”, “manhattan”, “canberra”, “minkowski”, and “mahalanobis”. Beyond these standard choices, *paRao()* extends its functionality to include all distance calculations available in the *proxy::dist()* function as well as user defined distance functions.

To compute Rao’s index using an *α* value of 1, along with the Euclidean distance between pixel values, for the Stebbins NDVI set of imagery, we employ the following approach:

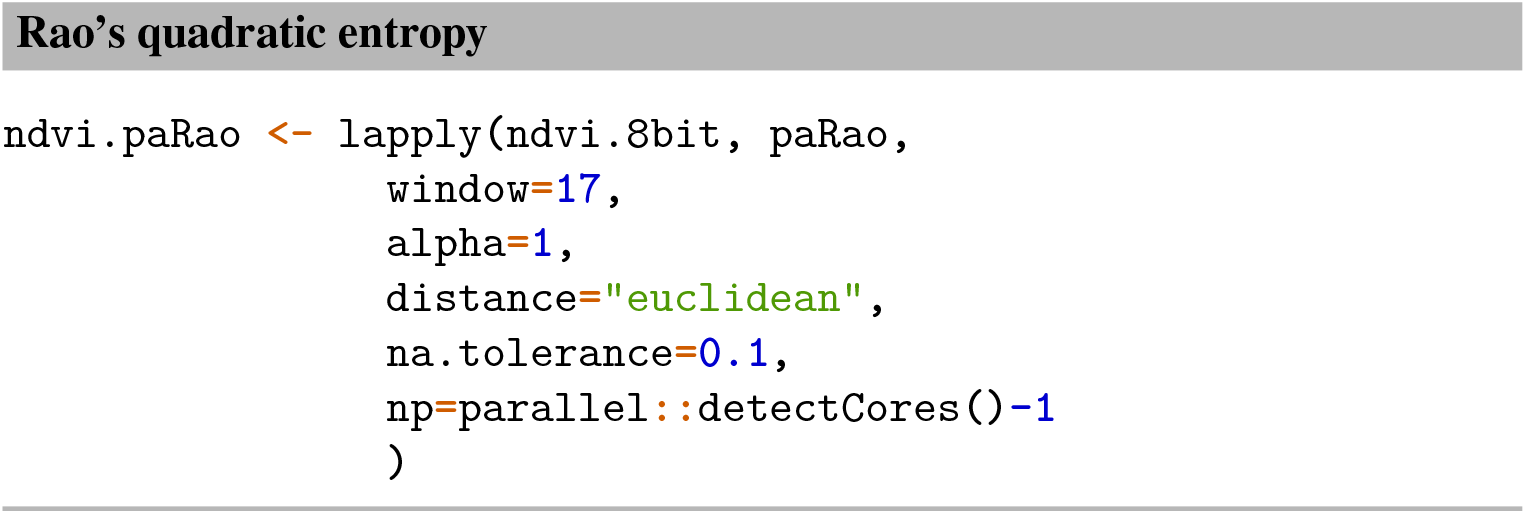

*paRao()* offers the alternative to calculate Rao’s index using defined areas instead of fixed moving windows. This functionality is accessible by supplying a polygonal vector layer through the *area* option, alongside the corresponding polygon identifier via the *field* option. In our upcoming example, we will demonstrate the computation of Rao’s index within specific areas of Stebbins Canyon, delineated by isopleths (contours of equal altitude), for NDVI observations spanning the three distinct years:

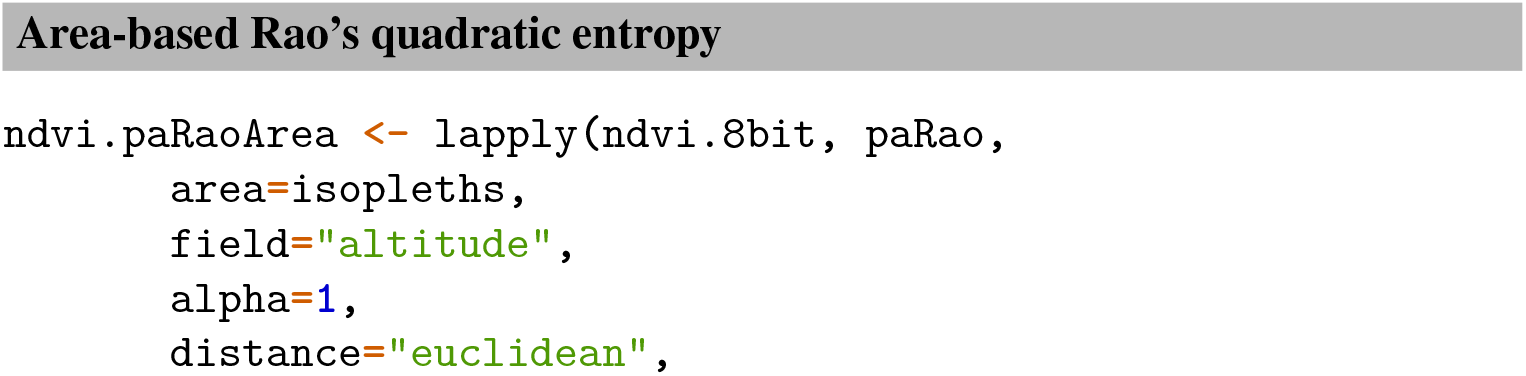
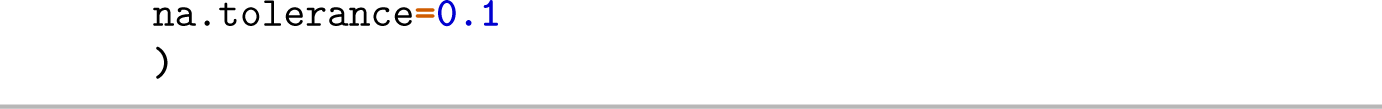

It is evident how Rao’s index varied considerably spatially in the 2016 imagery, which was taken less than two months after the Wragg wildfire (Figure 6). In the following years, the landscape underwent a strong homogenisation due to a decrease in Rao’s entropy index for formerly spectrally diverse habitats located at the bottom of the valley.

**Fig. 6.**
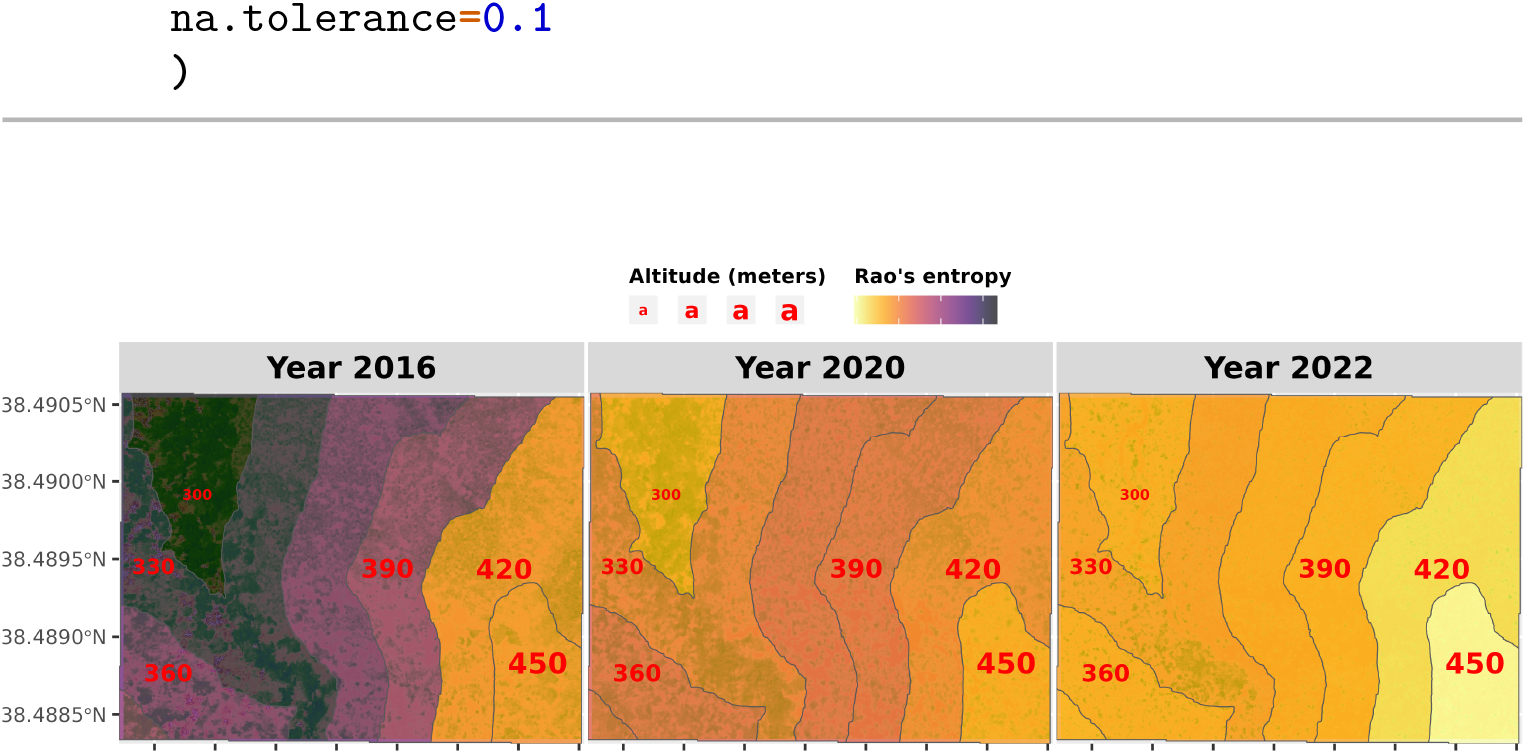
Overview of parametric Rao’s index calculated using areas defined by isopleths for NDVI in different years. Darker values show areas where pixel distances and thus spectral heterogeneity is higher.

### 3.1 Multidimensional Rao’s quadratic entropy

The extended version of the parametric Rao index implemented in *rasterdiv* facilitates the incorporation of multiple raster layers or numerical matrices into the diversity index computation. This approach calculates the distance matrix across the *n* raster layers within a multi-dimensional framework using the distance measures previously discussed. To utilise this methodology, it is essential to specify the option *method=“multidimension”* and to provide a minimum of two single-layer *spatRasters* either as a list or a vector, or a multi-layer *spatRaster*. In scenarios where a multi-layer *spatRaster* is provided, the *paRao()* function will automatically incorporate all layers in the index calculation.

In this example, we demonstrate the multi-dimensional application of *paRao()* by using it on the red and near-infrared bands (supplied as a multilayer spatRaster) from a sub-selection of the 2016 Stebbins Cold Canyon imagery. Thus, here we focus on the single spectral bands that are crucial for assessing vegetation dynamics, rather than on their combination to derive NDVI (Figure 7).

**Fig. 7.**
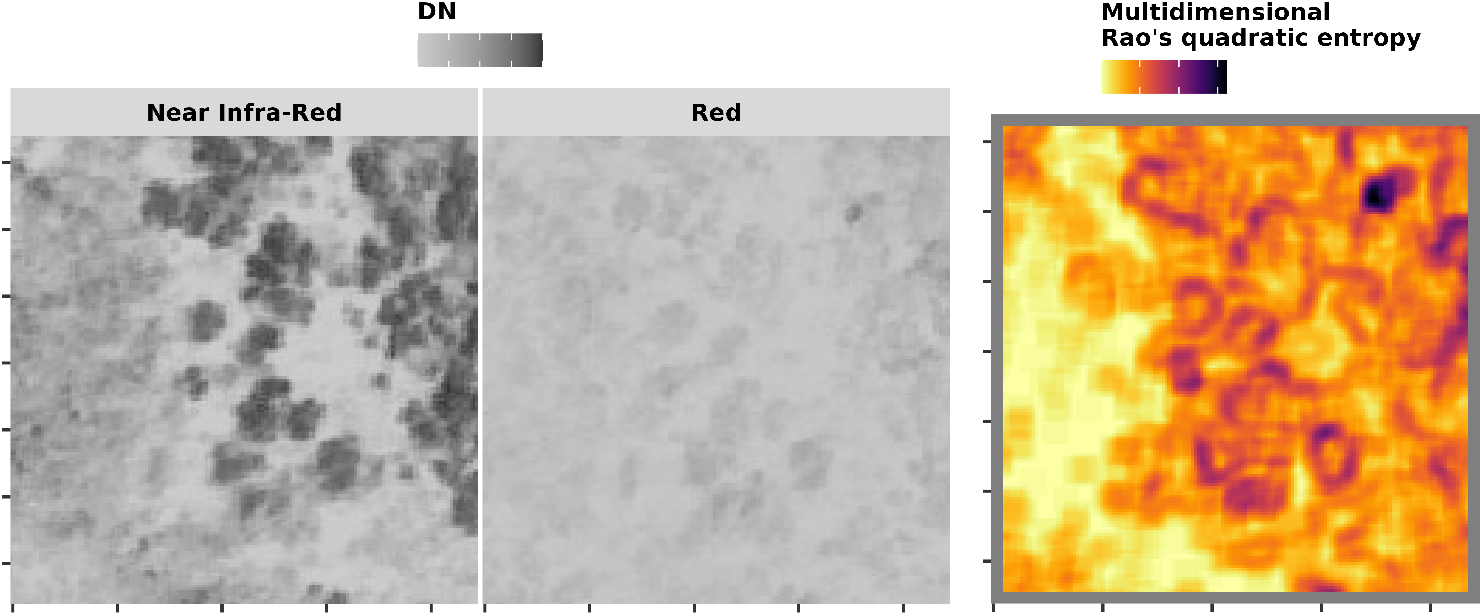
Overview of parametric Rao’s index calculated using areas defined by isopleths for NDVI in different years. Darker values show areas where pixel distances and thus spectral heterogeneity is higher.

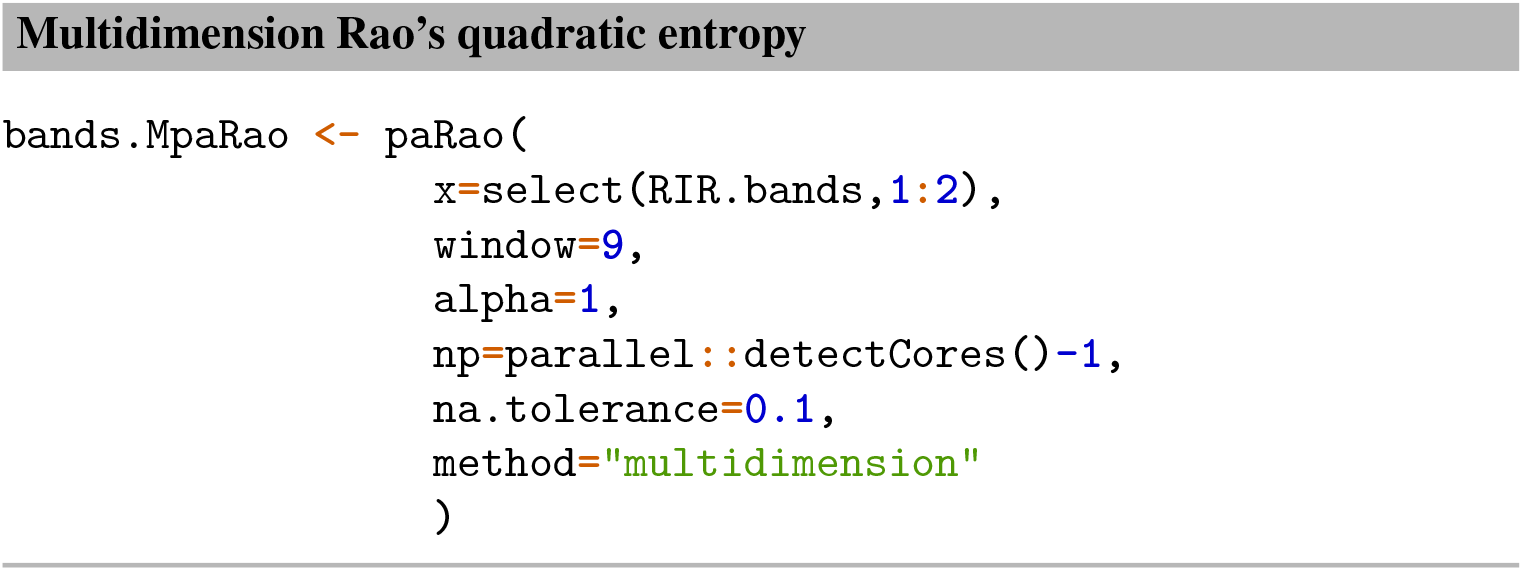

As for the other indexes in *rasterdiv*, multidimension *paRao()* allows for multiple moving window (*window*), *α* values (*alpha*), distance metrics (*dist m*) and simplification of the numerical matrices (*simplify*). Moreover, the option *rescale* can be used in multidimensional *paRao()* to centre and scale the values of each layer provided for calculation.

### 3.2 Rao’s Accumulation function

The *rasterdiv* package also includes the *AUCRao()* function, which calculates the Area Under the Curve (AUC) for the Rao’s index across a range of alpha values. This is especially valuable in exploratory data analysis where the choice of alpha for the Rao’s index may not be immediately apparent. By assessing the AUC, researchers can gain insights into the overall trend of diversity across different scales of emphasis on spectral classes abundance and distance [6].

The *AUCRao()* function is based on the *paRao()* function, meaning it supports all the arguments that *paRao()* does. This provides flexibility in handling different types of input data and distance measures. For instance, in the following example, the *AUCRao()* function is applied to integrate the Rao’s index over alpha values from 1 to 5, allowing us to compare the cumulative effect of these alpha parameters on the diversity index outcome.

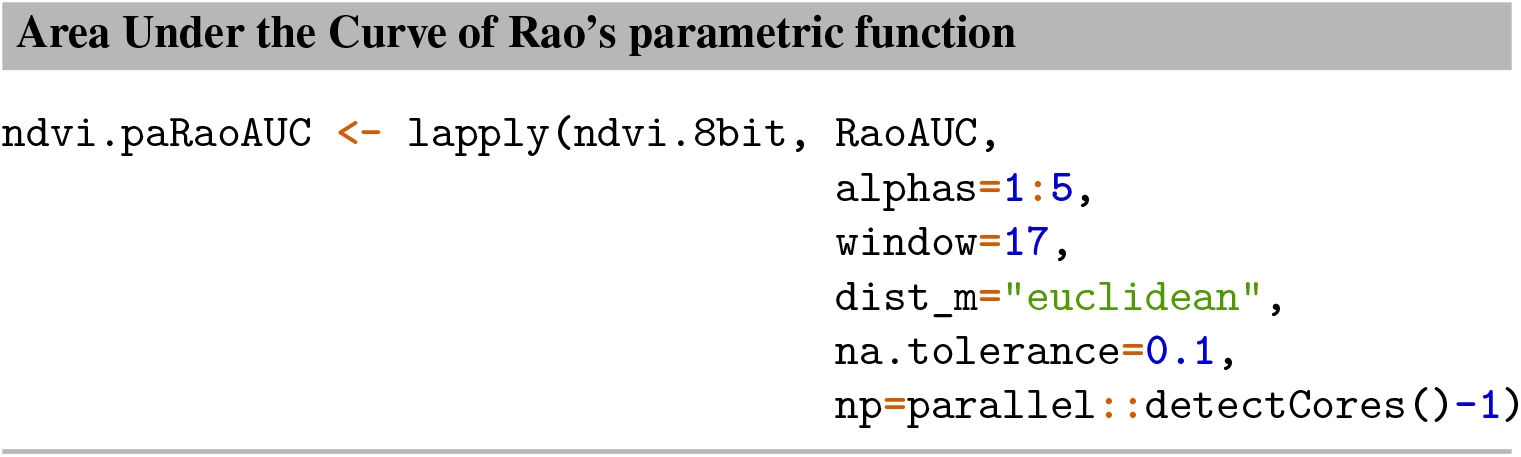

## 4 Helical Plots to visualise time series variability

Remote sensing data play a crucial role in monitoring and visualising temporal trends in various ecological and environmental contexts and spatial scales. These datasets, rich in spatial and temporal information, are pivotal in discerning subtle yet significant changes occurring over time. Traditional visualisation methods, while offering simplicity and clarity, often compress or overlook nuanced details that are vital for a comprehensive understanding of ecological dynamics [2].

In contexts where the complex interplay of various factors is common, more sophisticated visualisation techniques are required. One innovative approach is the use of “Helical Graphs” (H-graphs), developed by statistician Danny Dorling and illustrator Kirsten McClure. These graphs offer a unique visualisation method by intertwining the magnitude and directional trends of a metric with its temporal progression [3]. H-graphs stand out for their ability to convey complex data narratives, blending quantitative changes with their qualitative aspects, thus providing a more holistic view of the ecological changes observed. This approach not only enhances the interpretability of remote sensing data but also opens new avenues for communicating complex environmental transformations in a more intuitive and insightful manner [4].

In the *rasterdiv* package, Helical Graphs (H-graphs) are implemented via two distinct functions: *heliPrep()* and *heliPlot()*. The *heliPrep()* function is designed to calculate the rate of change between time-series values. It then applies a two-sided moving average linear filter to both the original values and their calculated rate of change. This function accepts three arguments: *values* for the time series of numeric values, *dates* for the corresponding chronological data, and *window_size* for determining the size of the linear filter.

The *heliPlot()* function is then responsible for visualising the dataframe generated by *heliPrep()*. It does so by leveraging B-splines within the *ggplot2* framework. This function, supplied with the data argument, enables the separation of input data into distinct helical lines when the *group* argument is utilised. Additionally, *heliPlot()* is versatile, allowing the incorporation of various *ggplot()* arguments using the + syntax from *ggplot2*, thus offering extensive customisation and flexibility in data visualization.

To show how to build H-plot with *rasterdiv* we will start from a dataframe presenting a multi-year time series of remotely sensed weekly NDVI values from a patch of old growth beech forest from central Italy. This plant community represents a relatively stable vegetation dynamics typical of broadleaved mountain forests. The dataframe *NDVI*.*ts* has three column, *ndvi* is the average weekly NDVI, *dates* is the date corresponding to the first day of each weekly NDVI value.

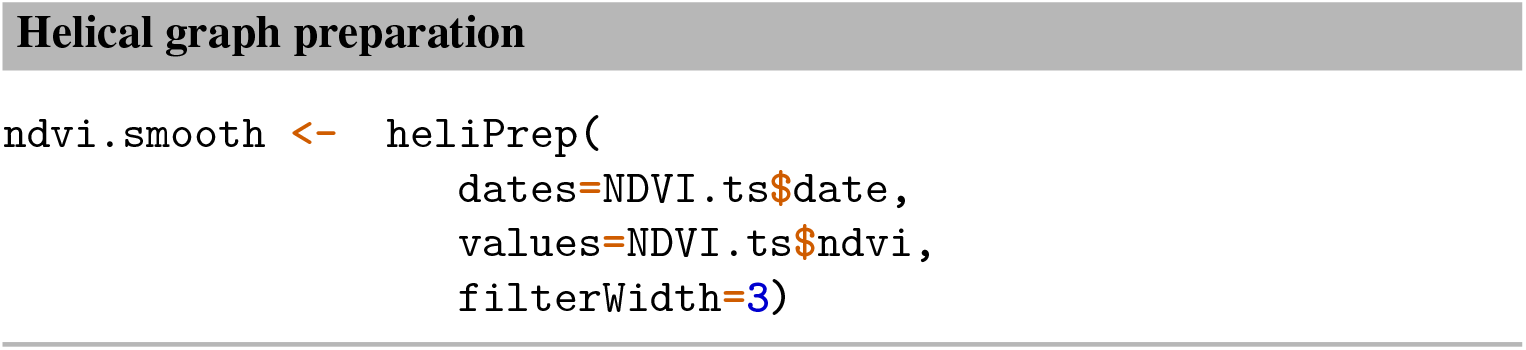

The output of *heliPrep(), ndvi*.*smooth*, is a dataframe with three columns, the averaged values, their rate of change and the corresponding date. We then use *heliPlot()* to visualise the H-plot:

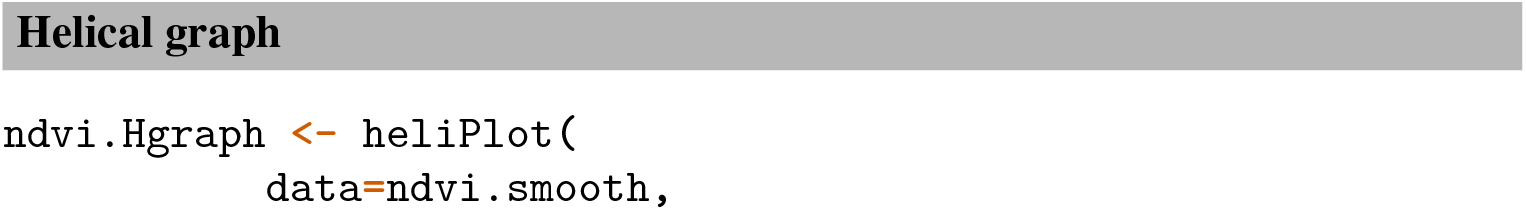
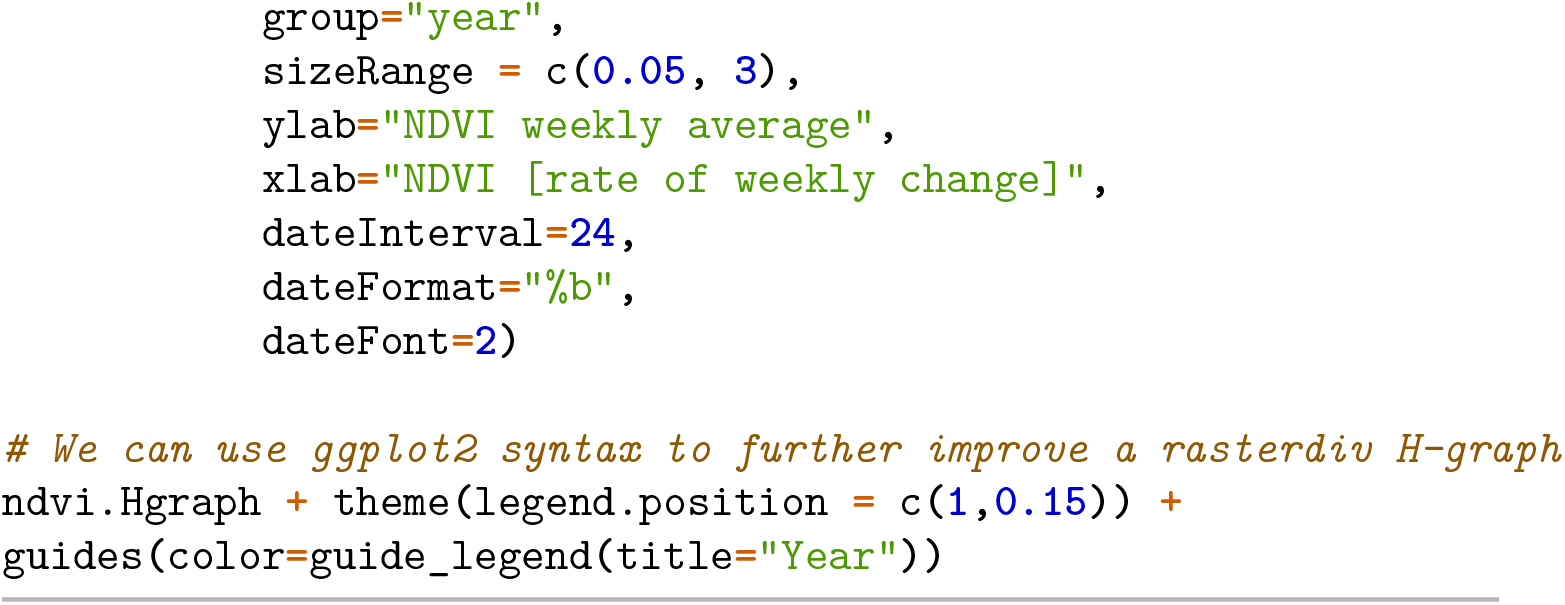

The H-plot provides a multi-dimensional view of vegetation dynamics, capturing both the magnitude of NDVI values over time and the rate at which these values change, thereby offering a comprehensive picture of vegetative health and seasonal changes in the beech forest (Figure 8). Lines circling upwards toward the ‘June’ marker denote increasing NDVI values, which typically correspond to growing biomass during the spring and early summer months. The closeness of these lines suggests that the annual vegetative patterns are consistent, with similar growth and senescence rates each year. This is expected for old-growth forests, which tend to have stable yearly cycles. The horizontal axis shows how quickly the NDVI values rise or fall. A steeper slope on this plot would indicate a more rapid change, such as during the onset of spring or the decline in autumn. The flatter parts of the curves suggest periods of little change, likely corresponding to the peak of summer or the dormancy of winter. If the different coloured lines show some divergence, this could indicate inter-annual variability in maximum NDVI values or in the timing of the vegetative peak. For instance, if one year’s line peaks later than others, it may suggest delayed spring growth, potentially due to climatic conditions. 2021 experienced comparatively lower levels of vegetation greenness for the beech forest patch, potentially as a consequence of an unusually cold and wet spring, which then gave way to a heatwave during the months of July and August. These climatic fluctuations may have stressed the vegetation, as evidenced by the NDVI’s reduced values. Conversely, the year 2018 is characterised by notably higher NDVI values in the spring, aligning with the fact that 2018 ranks as the second warmest year on record for Italy [5]. The warmer conditions likely promoted an earlier and more vigorous onset of the growing season, leading to heightened vegetative growth and greenness.

**Fig. 8.**
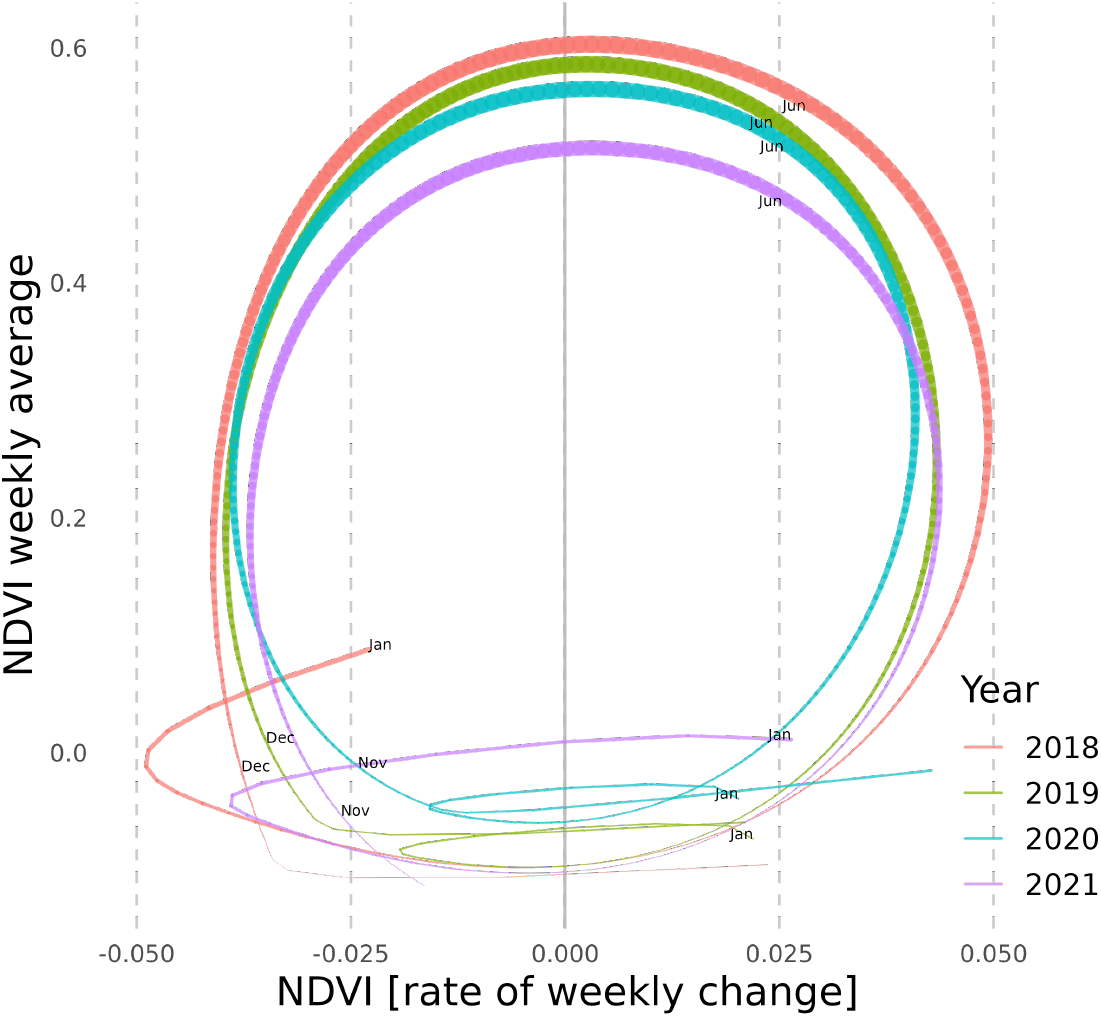
Helical plots of a multi-year weekly NDVI times series from a old growth beech forest in central Italy.

The utility of H-graphs extends beyond the visualisation of simple NDVI time series; they can effectively represent the temporal evolution of various diversity indices calculated through the *rasterdiv* package. By capturing the dynamic nature of ecological data, H-graphs offer a powerful tool for illustrating trends and detecting anomalies in time series data of ecological diversity.

## References

1. Rocchini, Duccio, Matteo Marcantonio, Daniele Da Re, Giovanni Bacaro, Enrico Feoli, Giles M. Foody, Reinhard Furrer, et al. 2021. “From Zero to Infinity: Minimum to Maximum Diversity of the Planet by Spatio-Parametric Rao’s Quadratic Entropy.” Global Ecology and Biogeography 30 (5): 1153–62. $10.1111/geb.13270$.

2. Cooksey, Ray W. 2020. “Descriptive Statistics for Summarising Data.” In Illustrating Statistical Procedures: Finding Meaning in Quantitative Data, edited by Ray W. Cooksey, 61–139. Singapore: Springer. $10.1007/978-981-15-2537-7_5$.

3. Dorling, Danny. 2020. Slowdown: The End of the Great Acceleration—and Why It’s Good for the Planet, the Economy, and Our Lives. Yale University Press.

4. Thouverai, Elisa, Matteo Marcantonio, Emanuela Cosma, Francesca Bottegoni, Roberto Cazzolla Gatti, Luisa Conti, Michele Di Musciano, et al. 2023. “Helical Graphs to Visualize the NDVI Temporal Variation of Forest Vegetation in an Open Source Space.” Ecological Informatics 74 (May): 101956. $10.1016/j.ecoinf.2022.101956$.

5. Istituto di Scienza dell’atmosfera e del clima. 2018. “2018 Warmest Year for Italy since 1800”. Accessed November 24, 2023. https://www.isac.cnr.it/en/news/2018-warmest-year-italy-1800.

6. Thouverai, Elisa, Matteo Marcantonio, Jonathan Lenoir, Mariasole Galfré, Elisa Marchetto, Giovanni Bacaro, Roberto Cazzolla Gatti, et al. 2022. “Integrals of Life: Tracking Ecosystem Spatial Heterogeneity from Space through the Area under the Curve of the Parametric Rao’s Q Index.” Ecological Complexity 52 (December): 101029. 10.1016/j.ecocom.2023.101029.

7. Rocchini, Duccio, Elisa Thouverai, Matteo Marcantonio, Martina Iannacito, Daniele Da Re, Michele Torresani, Giovanni Bacaro, et al. 2021. “Rasterdiv—An Information Theory Tailored R Package for Measuring Ecosystem Heterogeneity from Space: To the Origin and Back.” Methods in Ecology and Evolution 12 (6): 1093–1102. 10.1111/2041-210X.13583.

8. Marcantonio Matteo, Duccio Rocchini, and Gianluigi Ottaviani. 2014. “Impact of Alien Species on Dune Systems: A Multifaceted Approach.” Biodiversity and Conservation, July, 1–24. 10.1007/s10531-014-0742-2.

9. Planet Team (2017). Planet Application Program Interface: In Space for Life on Earth. San Francisco, CA. https://api.planet.com.

10. Rao, C. Radhakrishna. 1982. “Diversity and Dissimilarity Coefficients: A Unified Approach.” Theoretical Population Biology 21 (1): 24–43. 10.1016/0040-5809(82)90004-1.

11. Rényi, Alfréd. 1961. “On Measures of Entropy and Information.” In Proceedings of the Fourth Berkeley Symposium on Mathematical Statistics and Probability, Volume 1: Contributions to the Theory of Statistics, 4.1:547–62. University of California Press.

12. Rocchini, Duccio, Niko Balkenhol, Gregory A. Carter, Giles M. Foody, Thomas W. Gillespie, Kate S. He, Salit Kark, et al. 2010. “Remotely Sensed Spectral Heterogeneity as a Proxy of Species Diversity: Recent Advances and Open Challenges.” Ecological Informatics, Special Issue on Advances of Ecological Remote Sensing Under Global Change, 5 (5): 318–29. 10.1016/j.ecoinf.2010.06.001.

13. Hill, M. O. 1973. “Diversity and Evenness: A Unifying Notation and Its Consequences.” Ecology 54 (2): 427–32. 10.2307/1934352.

14. Boltzmann, L. 1872. Weitere studien über das wärmegleichgewicht unter gasmolekälen. S. K. Akad. Wiss. Wein 66.

15. Gini, C. 1912. Gini, Corrado. 1912. Variabilità e mutabilità: contributo allo studio delle distribuzioni e delle relazioni statistiche. [Fasc. I.]. Tipogr. di P. Cuppini.

16. Shannon, C. E. 1948. A mathematical theory of communication. Bell Syst. Tech. J. 27: 379–423, 623-656.

17. Simpson, E. H. 1949. Measurement of diversity. Nature 163:688.

18. Rao, Murali, Y. Chen, B.C. Vemuri, and Fei Wang. 2004. “Cumulative Residual Entropy: A New Measure of Information.” IEEE Transactions on Information Theory 50 (6): 1220–28.

19. Pielou, E. C. 1966. “The Measurement of Diversity in Different Types of Biological Collections.” J. Theor. Biol. 13: 131–44.

20. Berger, Wolfgang H., and Frances L. Parker. 1970. “Diversity of Planktonic Foraminifera in Deep-Sea Sediments.” Science 168 (3937): 1345–47.

